# Inhibition of constitutive activity of the atypical chemokine receptor ACKR3 by the small-molecule inverse agonist VUF16840

**DOI:** 10.1101/2024.12.30.630720

**Authors:** Reggie Bosma, Desislava Nesheva, Merel Rijnsburger, Rick Riemens, Justyna M. Adamska, Max Meyrath, Simon Mobach, C. Maurice Buzink, Iwan J. P. de Esch, Martyna Szpakowska, Maikel Wijtmans, Andy Chevigne, Henry F. Vischer, Rob Leurs

## Abstract

The atypical chemokine receptor 3 (ACKR3) has emerged as a promising drug target for the treatment of cancer, cardiovascular, and autoimmune diseases. In this study, we present the pharmacological characterization of VUF16840, the first small-molecule inverse agonist of ACKR3. VUF16840 effectively displaces CXCL12 binding to ACKR3 and inhibits chemokine-induced β-arrestin2 recruitment in a concentration-dependent manner. Furthermore, VUF16840 stabilizes the inactive conformation of ACKR3, as demonstrated by its ability to suppress constitutive β-arrestin2 recruitment. This inverse agonism alters ACKR3 constitutive trafficking, leading to receptor enrichment at the plasma membrane and inhibition of intracellular CXCL12 uptake. Importantly, VUF16840 exhibits high selectivity for ACKR3 over a broad panel of human chemokine receptors.

These findings establish VUF16840 as a potent and selective ACKR3 inverse agonist capable of modulating constitutive and chemokine-induced signaling and internalization events. As such, VUF16840 represents a valuable pharmacological tool for exploring the molecular and translational roles of ACKR3 in both physiological and pathological contexts.

## Introduction

The atypical chemokine receptor 3 (ACKR3), formerly known as CXCR7, was initially identified as a second receptor for the CXC-chemokine ligand CXCL12, previously thought to exert its effects solely via the CXC chemokine receptor 4 (CXCR4)^1,2^. Since its discovery, ACKR3 has been extensively investigated for its pathological roles, including its contributions to tumor progression and anti-tumor therapy resistance^3^. Additionally, ACKR3 has been implicated in cardiovascular diseases^4,5,6^, HIV entry into human host cells^7^ and autoimmune disorders^3^ underscoring its potential as a viable therapeutic target.

ACKR3 is a member of the G protein-coupled receptor (GPCR) superfamily and binds the chemokines CXCL11 and CXCL12 with sub-nanomolar affinities.^1,2^ More recently, ACKR3 has been shown to bind a number of opioid peptides^8^ suggesting a novel role for ACKR3 in pain perception. Unlike typical GPCRs, ACKR3 does not activate G proteins upon ligand binding.^2,9,10^ However, ligand binding increases ACKR3 interaction with β-arrestins, a process commonly used as a functional readout for ACKR3 activation.^8–12^ The recruitment of these intracellular regulatory proteins upon GPCR activation translates to diverse cellular responses for numerous GPCRs.^13, 14^ Upon chemokine binding, ACKR3 rapidly internalizes from the cell membrane, scavenging bound chemokines from the extracellular milieu for intracellular degradation^1,10,12^. This scavenging activity modulates the extracellular levels of CXCL11 and CXCL12, which are critical for directional cell migration mediated by CXCR3 and CXCR4, respectively, thereby influencing e.g. cancer metastasis and immune cell migration in e.g. multiple sclerosis (MS).^3,15^

The scavenging function of ACKR3 can be disrupted by ligands that inhibit chemokine binding, regardless of whether they are agonists or antagonists. Consequently, controversy has arisen regarding the classification of ACKR3 ligands. For instance, compounds such as CCX754, CCX662, CCX733, and CCX771, often described as inhibitors or antagonists of ACKR3,^15–19^ are capable of stimulating ACKR3-mediated β-arrestin2 recruitment.^12,16, 20^ In fact, to date most small molecules reported in the literature or natural compounds like conolidine^19,21,22^ induce β-arrestin recruitment to ACKR3. Accordingly, small molecule ACKR3 agonists have been widely used to probe its scavenging function, showing efficacy in various cancer models,^2,3,15,17,20^ while inhibition of ACKR3 activity remains underexplored.

Following up on previous work on small molecule ACKR3 agonists^21,23^, we launched a research program aimed at identifying ACKR3 antagonists. At the start of this project, potent ACKR3 antagonists were not described in the scientific literature. We therefore performed an analysis of patent literature, as several companies had applied for patent protection of various chemical scaffolds acting on ACKR3, but often referred to those as CXCR7 or ACKR3 modulators^19,21,24^. In this search, we identified a series of ligands described in patent application WO201819929 of Idorsia Pharmaceuticals^25^ and claimed to antagonize ACKR3. One of the isooxazole-based ligands, VUF16840 (**Fig. 1A**), was synthesized in-house and its detailed pharmacological characterization is described in this study. Notably, VUF16840 acts as an inverse agonist, suppressing constitutive signaling of ACKR3. This inverse agonism was observed in assays measuring the recruitment of β-arrestin1/2 or GRK2, as well as in assays related to receptor internalization dynamics, leading to reduced constitutive endosomal translocation, enhanced cell surface localization and reduced cellular CXCL12 uptake. Interestingly, VUF16840 inhibition with ACKR3 is non-surmountable by chemokines, resulting in potent inhibition even at high CXCL11 or CXCL12 concentrations. Moreover, VUF16840 demonstrates high selectivity for ACKR3 over other chemokine receptors, making it a valuable tool for investigating the role of ACKR3 constitutive activity and the therapeutic potential of ACKR3 antagonism in diverse biological contexts. During the course of our study, a close analog, ACT-1004-1239, was disclosed as an ACKR3 antagonist in the literature by Idorsia Pharmaceuticals^26, 27^ and presented as a clinical candidate for the treatment of MS^28–30^, making the present results relevant for the understanding of the potential therapeutic effect of ACKR3 modulation in MS.

**Figure 1.**
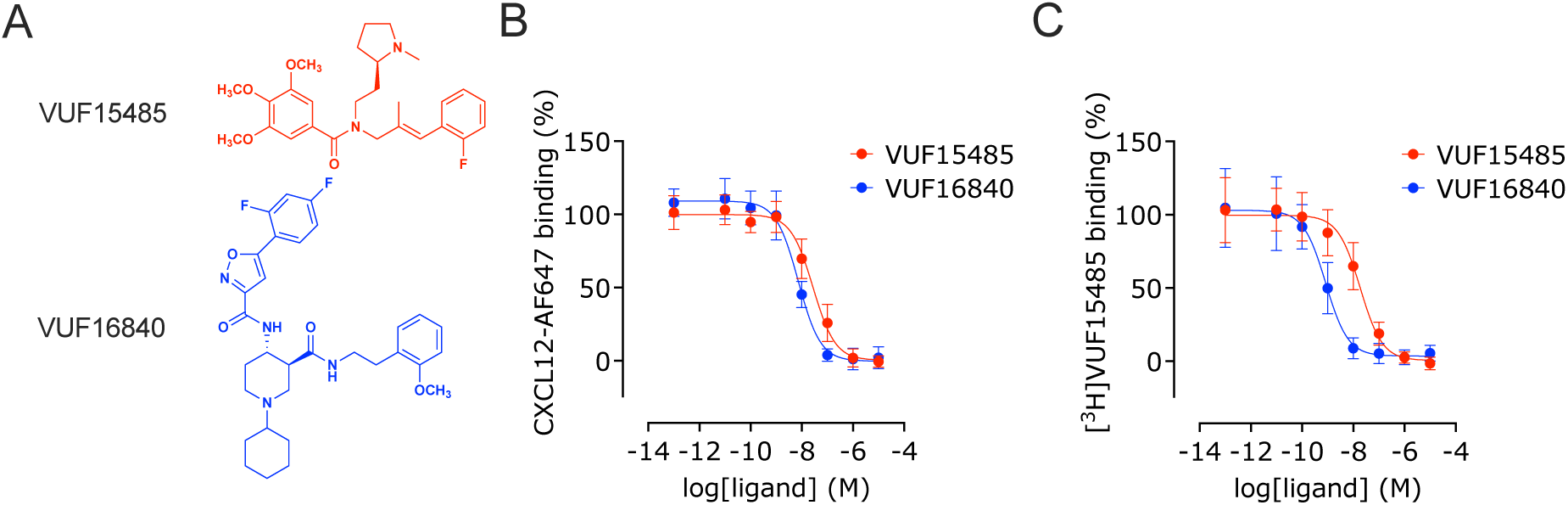
Concentration-dependent displacement of binding of labeled agonists to ACKR3. (**A**) Structures of displacers VUF 15485 and VUF16840. HEK293T membranes transiently expressing either NLuc-hACKR3 or HA-hACKR3 were used to measure (**B**) the concentration dependent displacement of the binding of fluorescently labeled CXCL12-AF647 or (**C**) the displacement of the binding of radioligand [^3^H]VUF15485 by unlabeled VUF15485 and VUF16840. Percentage of residual probe binding was normalized and the depicted data are the mean ± SD of ≥ 3 experiments with triplicate measurements per experiment.

## Materials and methods

### Materials

HBSS (with Ca^2+^ and Mg^2+^), Dulbecco’s Modified Eagle’s Medium (DMEM; high glucose), 0.05% trypsin solution and penicillin/streptomycin solution was purchased from Thermo Scientific (Waltham, United States). Materials for radioligand binding, GF/C filter plates and Microscint-O, were purchased from Perkin Elmer (Waltham, United States). Fetal bovine serum (FBS) was obtained from Bodinco (Alkmaar, the Netherlands). Linear 25 kDa polyethylenimine (PEI) was purchased from Polysciences (Warrington, United States). Bovine serum albumin (BSA) was obtained from Melford (Ipswich, United Kingdom). NanoGlo and coelentrazine-h substrates were purchased from Promega (Madison, United States). Resazurin is obtained from Merck (Kenilworth, United States). 96-well plates were obtained from Greiner Bio-one (Kremsmünster, Austria). White low volume 384 well plates were bought from Corning (Corning, United States).

VUF15485 and VUF16840 were synthesized and characterized in-house (**Fig S1-S4**). Human recombinant CXCL11 (#CN-13), CXCL12 (#CN-11) and fluorescently labeled CXCL12-A647 (#CAF-11) were purchased from Almac (Craigavon, United Kingdom) or Protein Foundry (Milwaukee, United States). All used chemicals were of analytical grade purity.

### Plasmids

The NLuc-ACKR3 plasmid was a kind gift from Prof Hill (Nottingham University, Nottingham, United Kingdom)^31^, Venus-Rab5a was a kind gift from Dr. Lambert (Georgia Health Sciences University, Augusta, GA, USA)^32^ and β-arrestin2-LgBiT/pBiT1.1-C[TK/LgBiT] and SmBiT-clathrinA/pBiT2.1-N[TK/SmBiT] plasmids were kindly provided by Dr. Seong (Korea University, Seoul, Republic of Korea).^33^ The mVenus -CAAX and the hACKR3-NLuc constructs have been described by Zarca et al^23^. The sequence of the used ACKR3 constructs corresponds to NCBI database entry NM_020311.3. Radioligand binding studies utilised a mammalian expression vector pcDEF_3_ encoding ACKR3, N-terminally fused to an HA-tag and C-terminally fused to a triple Alanine tag (NotI). This construct was the template for the C-terminal ACKR3 fusion with the luminescent Rluc8 protein (NCBI: EF446136.1). The CXCR4-Rluc8 fusion protein corresponds to CXCR4 isoform b (NCBI: NM_003467.2) fused at the C-terminus with a triple Alanine linker (NotI) before the Rluc8 protein.

For BRET experiments, several BRET acceptor constructs were used that consist of the pcDEF_3_ vector with either GRK2 (NCBI: NM_174710.2), β-arrestin1 (NCBI: NM_174243.3) or β- arrestin2 (NCBI: XM_005220181.4) fused at the C-terminus with a SpeI/NotI linker to mVenus (NCBI: KP666136.1), as previously described.^34^

For NanoBiT-complementation experiments, the HA-tagged ACKR3 was C-terminally fused to a SmBiT separated by a TSSGSSGGGGSGGGGSS-linker in pcDEF_3_, as previously described.^35^ The LgBiT-β-arrestin2 construct was generated by flanking the open reading frame of β- arrestin2 with SacI and XbaI restriction sites at the N- and C-termini, respectively, by PCR using the aforementioned β-arrestin2-mVenus as template. The resulting β-arrestin2 fragment was subcloned in the pBiT1.1-N[TK/LgBiT] vector (kindly provided by Promega) using the indicated restrictions enzymes. The NanoBiT constructs that were used for screening the pharmacological activity of VUF16840 on the panel of human chemokine receptors were described before.^8^

### Cell culture

HEK293T cells were maintained in DMEM, supplemented with 10% fetal bovine serum and 1x penicillin/streptomycin. Cells were grown in 10 cm dishes and kept in a humidified atmosphere with 5% CO_2_ at a temperature of 37°C. When cells reached a near confluent monolayer (∼85%), or in preparation of further experimentation, cells were briefly incubated with a 0.05% trypsin solution and subsequently collected in culture medium. Cell numbers were quantified using a TC20 cell counter from BioRad (Hercules, United States) and subsequently seeded at the appropriate density for propagation or experimentation.

### Cell transfection

For 1·10^6^ HEK293T cells, a transfection mix consisting of 2 µg plasmid DNA, 12 µg linear polyethylenimine (PEI) in a total volume of 200 µL 150 mM NaCl was made and thoroughly vortexed. The transfection mix was incubated for 15 min and consequently added to a cell suspension with a final concentration of 3·10^5^ cells per mL. For functional studies this cell suspension was immediately transferred to a poly-L-lysine-coated white 96-well plate, with 3·10^4^ cells per well. For BRET experiments, the 2 µg plasmid DNA of the transfection mix consisted of 0.4 µg Rluc8-fused chemokine receptors (ACKR3-Rluc8 or CXCR4-Rluc8) and 1.6 µg of one of the mVenus-fusion proteins (β-arrestin1-mVenus, β-arrestin2-mVenus, GRK2-mVenus, Rab5a-mVenus). For NanoBiT-complementation experiments detecting β-arrestin2 recruitment to the ACKR3, the 2 µg DNA of the transfection mix consisted of 0.4 µg ACKR3-SmBiT, 0.6 µg β-arrestin2-LgBiT and 1 µg pcDEF_3_. Finally, to detect the interaction between β-arrestin2 and Clatherin A using NanoBiT, the 2 µg DNA of the transfection mix consisted of 0.4 µg ACKR3, 0.6 µg β-arrestin2-LgBiT, 0.6 µg SmBiT-Clathrin A and 0.4 µg pcDEF_3_.

### Membrane preparations

Membranes of HEK293T cells that transiently expressed the NLuc-hACKR3 or HA-hACKR3 were produced as previously described^23^. Briefly, 2 million HEK293T cells were seeded per 10 cm^2^ dish and were transfected the following day using a DNA/PEI-mix containing 0.25 µg plasmid DNA encoding for the ACKR3, 4.75 µg pcDEF3 plasmid DNA and 30µg linear PEI. Two days after transfection, cells were collected in PBS [137 mM NaCl, 2.57 mM KCl, 1.5 mM KH_2_PO_4_, 8 mM Na_2_HPO_4_] and pelleted by centrifugation. Cell pellets were resuspended in ice-cold membrane buffer [15 mM Tris, 0.3 mM EDTA, 2 mM MgCl_2_, pH 7.4 at 4°C] and dounce-homogenized on ice by plunging the pestle 10 times with 1500 rpm. Cell homogenates were subsequently subjected to two freeze-thaw cycles using liquid nitrogen. Next, the cell homogenates were centrifuged at 40,000 g and pellets were resuspended in Tris-sucrose buffer [20 mM Tris, 250 mM sucrose, pH 7.4 at 4°C]. Finally, the membrane samples were homogenized using a 23-gauge needle, snap-frozen with liquid nitrogen and stored until further experimentation at -80 °C.

### NanoBRET ligand binding assay

NanoBRET binding was measured in triplicate on white low-volume 384-well plates and started by combining 0.3 nM fluorescently labelled CXCL12 (CXCL12-AF647), increasing concentrations of VUF16840 or VUF15485 (10^-5^ M – 10^-11^ M) with 35 ng membranes (protein content) per well of HEK293T cells expressing NLuc-ACKR3. All dilutions were prepared in HBSS supplemented with 0.2% BSA. After assembling the binding reaction, the plate was flash-centrifuged at 100g. Binding reactions were then incubated for 1 hour after which NanoGlo substrate was added (310 times diluted from stock concentration) to a final volume of 13.5 µL. Total light intensity was measured at 460 nm wavelength with 80 nm bandwidth and separately for wavelengths ≥610 nm using the PHERAstar-FSX (BMG Labtech, Ortenberg, Germany) with a dual emission filter. The ratio of light intensities (>610 nm over 460 nm) is a measure for the relative binding of CXCL12-AF647 to the NLuc-ACKR3.

### [^3^H]VUF15485 radioligand binding assay

Radioligand binding studies were performed as described earlier^23^. Membranes expressing HA-ACKR3 were incubated with 4 μg membrane protein/well and 2 nM [^3^H]VUF15485 and increasing concentrations VUF16840 or VUF15485, diluted in in HBSS supplemented with 0.2% BSA. All assays were performed in triplicate in a 96-well plate. Binding reactions were incubated for 2 hours at room temperature and then terminated by filtration over a PEI-coated GF/C filter using a cell harvester (Perkin Elmer) followed by three consecutive wash steps using ice-cold buffer [50 mM HEPES, 1.2 mM CaCl_2_, 5 mM MgCl_2_, 0.5 M NaCl, pH 7.4 at 4°C]. Filter-bound [^3^H]VUF15485 was measured by adding 25 µL Microscint-O per well to the dried GF/C-plate and radioactivity was quantified using the Wallac Microbeta counter (Perkin Elmer).

### Recruitment of signal transducers and receptor trafficking

Two days after transfection, medium was aspirated and cells were reconstituted in HBSS with 0.05% BSA, 5 µM coelenterazine-h (Rluc8 substrate) and various concentrations of CXCL11, CXCL12, VUF15485 and/or VUF16840 in triplicate. Cells were subsequently incubated at 37 °C for 20 min after which the total light intensity was measured at 475 nm and 535 nm (both with a bandwidth of 30 nm) using the PHERAstar-FSX with a dual emission filter. The ratio of light intensities (535 nm over 475 nm) is a measure for the relative abundance in proximity between the respective chemokine receptor and the respective mVenus-fused protein.

Ligand-induced receptor mobilisation to the plasma membrane was also monitored by bystander NanoBRET^36^. For this assay, 5 × 10^6^ HEK293T cells were seeded in 10-cm dishes and co-transfected with plasmids encoding ACKR3 C-terminally tagged with NLuc and mNeonGreen C-terminally tagged with the plasma membrane targeting sequence corresponding to the first 11 residues (MGCIKSKGKDS) of the human Lyn-kinase^37^. Twenty-four hours after transfection, cells were distributed into black 96-well plates (1 × 10^5^ cells per well) and treated with chemokines (100 nM). After 45-minute incubation at 37 °C, coelenterazine H (10 µM) was added and donor emission (460 nm) and acceptor emission (535 nm) were immediately measured on a GloMax Discover (Promega) plate reader.

### NanoBiT-complementation assay to measure β-arrestin1/2 or GRK2 recruitment

Two days following transfection (see above), medium was aspirated and cells were reconstituted in HBSS with 0.05% BSA and various concentrations of VUF15485 or VUF16840 in triplicate. Cells were incubated at 37 °C for 15 min after which NanoGlo substrate was added (310 times diluted from stock concentration). Cells were incubating at 37 °C for another 3 min and bioluminescent intensities were measured using the PHERAstar-FSX. Bioluminescence is a measure for the relative proximity of LgBiT- and SmBiT-tagged proteins and thus the occurrence of investigated protein-protein interaction^38^.

### CXCL12 uptake in ACKR3-overexpressing hCMEC/D3 cells

Human CMEC (D3) cells stably overexpressing ACKR3 and empty vector control were generated by retroviral transductions. The lentiviral expression vector encoding the human ACKR3 was created by cloning the ACKR3 gene (NM_020311.3 from GenScript) in the restriction sites Nhe1/Sma1 of the pLV-LifeAct (a kind gift of Dr. Niels Heemskerk). Lentiviral particles were packaged in HEK293T cells, cultured in DMEM containing 10% FCS, 1% penicillin/streptomycin (ThermoFisher scientific) at 37°C in a 5% CO_2_ incubator. Lentiviral expression and packaging vectors were introduced using calcium phosphate transfection. Supernatant containing virus was collected and virus was concentrated by centrifugation (Hettich swing-out rotor, 1500rpm), aliquoted (Amicon Ultra15, UFC910024 Merck) and stored at −80°C. For transduction of hCMEC/D3 endothelial cells, virus-containing supernatant was added dropwise to hCMEC/D3 cells in a 24 well-microplate. Virus supernatant was replaced with appropriate medium after 24 h incubation. Transduced cells were selected using puromycin (1:2000). For uptake assays, D3-ACKR3+ cells were grown to confluence in 96 wells plates in EG-2MV medium (Lonza; 1#CC3202). Cells were pre-treated for 30 minutes with 1, 10 or 100 nM VUF16840, where after 3 nM CXCL12-A647 (Protein Foundry) was added to the medium for 60 min. Supernatant was taken off, cells were washed 1x with medium and CXCL12-A647 signal in cells was measured using a BioTek Cytation microplate reader (Agilent, Santa Clara, USA). For immunocytochemistry (ICC), D3-ACKR3+ cells were cultured and stimulated as above on 8 well µ-slides (Ibidi systems), where after they were fixed with 4% PFA for 10 minutes. Cells were washed with PBS and permeabilized for 5 min in PBS+0.05% Triton-X after they were blocked with 10% goat serum for 60 min at RT. Rhodamine phalloidin (Molecular Probes, #R415; 1:400) was added for 1 hour, after which cells were washed, incubated with Hoechst (1:1000) for 5 minutes, washed and covered with mowiol on coverslips. Images were taken using a confocal microscope at 40X magnification.

### Data analysis

The sigmoidal dose-dependent displacement of binding of the probes [^3^H]VUF15485 and CXCL12-A647 by unlabelled ligands were analysed using Graphpad Prism 8, by fitting the data to a one-site binding model to determine the pIC_50_. Binding affinities (K_i_ values) for VUF15485 and VUF16840 were determined using the Cheng-Prusoff equation.^39^ Mean pIC_50_ values and pK_i_ values were determined by averaging fitted values from individual experiments. Graphs depict the pooled data of all experiments. Similarly, sigmoidal, ligand-induced concentration-dependent activation or inhibition of the proximity between ACKR3 and the protein of interest was analysed using Graphpad Prism 8, by fitting the data to a three-parameter dose-response model to determine pEC_50_ /pIC_50_ values. Mean pEC_50_ values and pIC_50_ values were determined by averaging fitted values from individual experiments. Graphs depict the pooled data of all experiments, for which values are normalized, per experiment, to the fold difference compared to basal signal. CXCL12 uptake and uptake inhibition by endothelial cells was analyzed with Graphpad Prism 10.2.0 using one-way ANOVA.

## Results

### VUF16840 binds ACKR3 with nanomolar affinity

The ACKR3 ligand VUF16840 was evaluated for its binding to hACKR3 in different competition binding assay formats (**Fig. 1**). Binding of fluorescent Alexa Fluor647-labeled CXCL12 (CXCL12-AF647) to membranes expressing N-terminally NLuc-tagged hACKR3 was quantified by a homogeneous NanoBRET binding assay^40^, whereas binding of [^3^H]VUF15485 to HA-tagged hACKR3 expressing membranes was determined after filtration of incubation mixtures and scintillation counting^23^. Co-incubation of CXCL12-A647 with NLuc-tagged hACKR3 expressing membranes and increasing concentrations of VUF16840 or the high-affinity small-molecule ACKR3 agonist VUF15485^23^ resulted in a concentration-dependent decrease in CXCL12-A647 binding to NLuc-hACKR3 (**Fig. 1A**). Both unlabeled ligands displaced CXCL12-A647 binding to NLuc-hACKR3 to similar levels with high inhibitory potencies; VUF15485 displaced CXCL12-A647 binding to NLuc-hACKR3 with a pIC_50_ value of 7.6 ± 0.1) (mean ± SD, n = 3; IC_50_ = 28 nM), whereas VUF16840 showed a pIC_50_ value of 8.2 ± 0.1 (mean ± SD, n = 3; IC_50_ = 7 nM). Both compounds were also potent in displacing the binding of [^3^H]VUF15485 from HA-hACKR3 expressing membranes. As shown in **Fig. 1B**, VUF16840 fully displaced [^3^H]VUF15485 binding from hACKR3 with a pIC_50_ of 9.1 ± 0.0 (mean ± SD, n = 13; IC_50_ = 0.7 nM). As previously shown (Zarca et al, 2024), VUF15485 potently displaced [^3^H]VUF15485 binding from the hACKR3 with a pIC_50_ of 7.7 ± 0.1 (mean ± SD, n = 18; IC_50_ = 20 nM.)

### VUF16840 inhibits constitutive β−arrestin2 recruitment to ACKR3

In the absence of G protein coupling, ACKR3 activation upon ligand binding is typically assessed by monitoring β−arrestin2 recruitment^36^. VUF16840 was therefore tested in a NanoBiT-based hACKR3-βarrestin2 recruitment assay (**Fig. 2**). As expected, addition of the ACKR3 agonist CXCL12 (0.14-100nM) to HEK293T cells transiently expressing hACKR3-SmBiT and β-arrestin2-LgBiT resulted in a rapid and concentration-dependent bioluminescence signal, indicative of successful NLuc complementation. Whilst low concentrations of CXCL12 produced a stable signal, higher agonist concentrations lead to a signal that gradually reduced over time (**Fig. 2A**), possibly due to increased ACKR3 internalization (vide infra).

**Figure 2.**
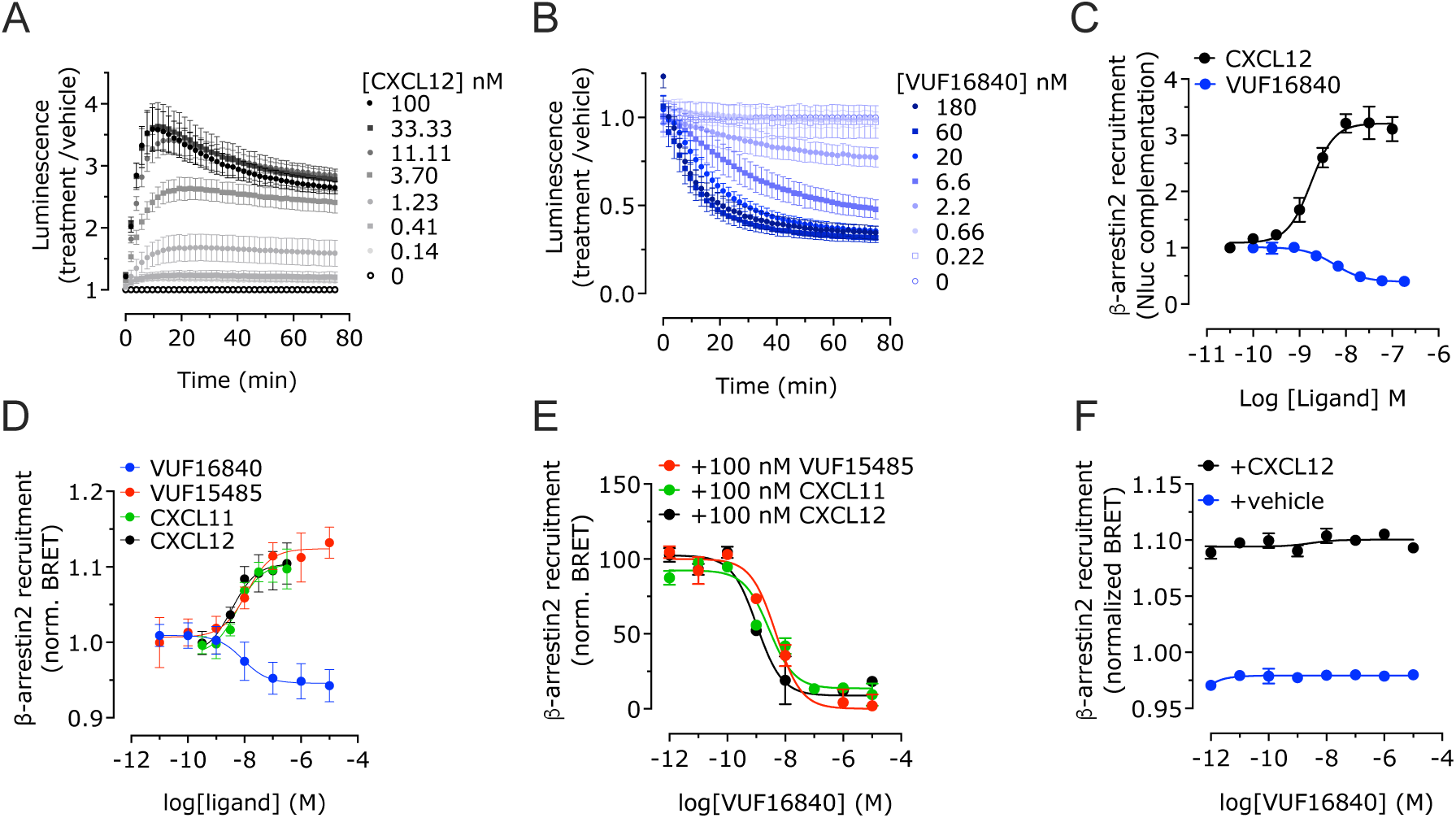
VUF16840 inhibits constitutive as well as agonist induced β-arrestin2 recruitment to ACKR3. (**A**) Transiently transfected HEK293T cells, expressing hACKR3-SmBiT and β-arrestin2-LgBiT, were stimulated with increasing concentrations CXCL12 or (**B**) VUF16840. Subsequently, β-arrestin2 recruitment to hACKR3 was detected by NanoBiT complementation. (**C**) Transfected HEK293T cells were stimulated for 30 min with increasing concentrations CXCL12 or VUF16840 and β-arrestin2 recruitment to the hACKR3 was detected by NanoBiT complementation. (**D**) Transfected HEK293T cells were stimulated with increasing concentrations VUF15485, VUF16840, CXCL12 or CXCL12 and β-arrestin2 recruitment to hACKR3 was detected by BRET. (**E**) β-arrestin2 recruitment to the ACKR3 was measured after stimulation with 100 nM of VUF15485, CXCL11 or CXCL12 in the presence of increasing concentrations VUF16840. (**F**) Transfected HEK293T cells were stimulated with increasing concentrations VUF16840 in the presence or absence of 100 nM CXCL12 and β-arrestin2 recruitment to the hCXCR4 was detected by BRET. Depicted data is normalized as fold-basal and represents the mean ± SD of ≥ 3 experiments with triplicate measurements per experiment.

In the same experimental set-up, VUF16840 did not lead to an increase in hACKR3 β-arrestin2 recruitment, but interestingly, the ligand time and concentration-dependently inhibited β-arrestin2 recruitment, suggesting constitutive coupling of hACKR3 to β-arrestin2 (**Fig. 2B**). Analysis of concentration-response curves generated after 30 min of incubation resulted in a pEC_50_ value of 8.8 ± 0.2 (mean ± SD, n = 3; EC_50_ = 1.6 nM) for the endogenous agonist CXCL12 and a pIC_50_ value of 8.2 ± 0.1 (mean ± SD, n = 3; IC_50_ = 6.3 nM) for the inverse agonist VUF16840 (**Fig. 2C**).

In an orthogonal BRET-based assay, HEK293T cells were transiently transfected with β-arrestin2-mVenus and hACKR3-Rluc8, allowing detection of β-arrestin2 recruitment to hACKR3 by BRET. As observed before^23^, the endogenous agonists CXCL11 and CXCL12 and the synthetic small molecule ACKR3 agonist VUF15485 induced concentration-dependent recruitment of β-arrestin2 to hACKR3 with nanomolar potencies (**Fig. 2D, Table S1**). In contrast, VUF16840 induced a concentration-dependent reduction of β-arrestin2 recruitment to hACKR3 with a pIC_50_ value of 8.0 ± 0.1 (mean ± S.D., n = 7; IC_50_ = 10 nM). The data obtained in this orthogonal assay corroborate the earlier findings of VUF16840 acting as an inverse agonist and hACKR3 showing constitutive coupling to β-arrestin2. The observed inverse agonism in a β-arrestin2 recruitment assay has so far not been reported for ACKR3 ligands and constitutive recruitment of a β-arrestin2 has so far only been incidentally reported for other GPCRs.^41–44^

In line with the aforementioned binding experiments with labelled CXCL12, VUF16840 competed with ACKR3 agonists in a β-arrestin2 recruitment assay (**Fig. 2E**). The inverse agonist exhibited a full concentration-dependent inhibition of ACKR3-β-arrestin2 interactions, induced by both CXCL11 and CXCL12 chemokines, as well as the small molecule agonist VUF15485 (100 nM) (**Fig. 2E**, **Table S1**). Yet, VUF16840 did not affect β-arrestin2 recruitment to the related CXCR4 receptor, neither in the presence nor absence of the shared, endogenous agonist CXCL12 (**Fig. 2F**). In a more detailed analysis (**Fig. 3**), ACKR3 expressing cells were pretreated with different concentrations of VUF16840 and concentration-response-curves of the ACKR3 ligands CXCL12 and CXCL11 were measured (Schild analysis). In this assay set-up, VUF16840 inhibited both CXCL12- and CXCL11-mediated β-arrestin2 recruitment in a non-surmountable fashion (**Fig. 3**). Whereas there is no actual shift in the EC_50_ values for the chemokines observed, the maximal response at high agonist concentrations is reduced concentration-dependently.

**Figure 3.**
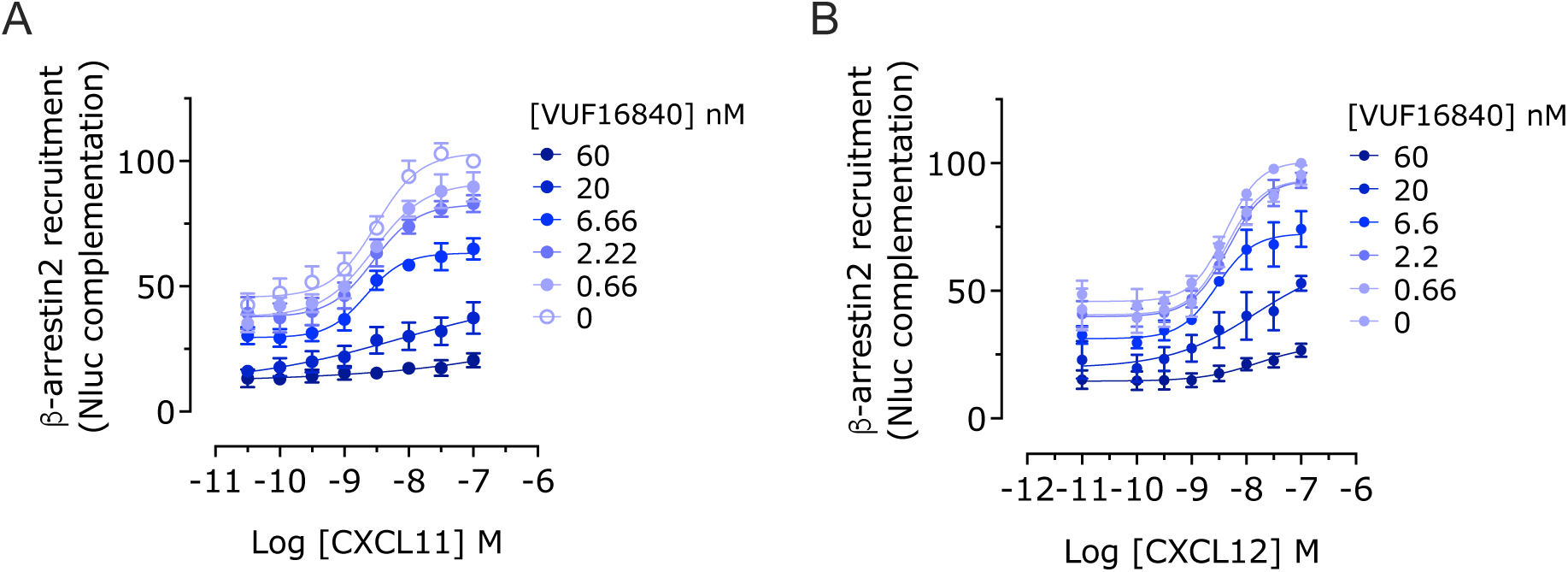
VUF16840 non-competitively inhibits agonist-induced β-arrestin2 recruitment to hACKR3. (**A**) Transiently transfected HEK293T cells, expressing hACKR3-SmBiT and β-arrestin2-LgBiT were stimulated with increasing concentrations of (**A**) CXCL11 or (**B**) CXCL12 following a 60 min pretreatment with buffer or various concentrations of the inverse agonist VUF16840. Next, the β-arrestin2 recruitment to the hACKR3 was detected by NLuc complementation.

### Chemokine receptor selectivity of VUF16840

To verify whether inverse agonism by VUF16840 is hACKR3 specific, the ligand was investigated for agonistic and antagonistic properties towards a panel of 19 human chemokine receptors (**Fig. 4**). To achieve this, HEK293T cells were transiently transfected with each chemokine receptor, C terminally tagged with the SmBiT NanoBiT sensor in combination with β-arrestin2, N-terminally fused to the LgBiT sensor. Transfected cells were stimulated with the respective endogenous agonist of each chemokine receptor, with a high concentration (1 μM) VUF16840 or with the combination of the two. In the agonist assay set-up VUF16840 stimulation only modulated β-arrestin2 recruitment to hACKR3 and hCCR3 (**Fig. 4A**). As expected, VUF16840 inhibited basal β-arrestin2 recruitment to hACKR3. Surprisingly, VUF16840 acted as an agonist for CCR3-mediated β-arrestin2 recruitment (**Fig. 4A**). Full concentration-response analysis (**Fig. SI5**) showed that VUF16840 was able to activate CCR3 with sub-micromolar potency (pEC_50_ = 6.7 ± 0.1; EC_50_ = 200 nM), whereas its endogenous ligand CCL13 activated CCR3 in our hands with almost equal potency (pEC_50_ = 7.2 ± 0.1, EC_50_ = 63 nM). Interestingly, VUF16840 activates CCR3 with higher efficacy (1.4-fold) than the endogenous agonist CCL13.

**Figure 4.**
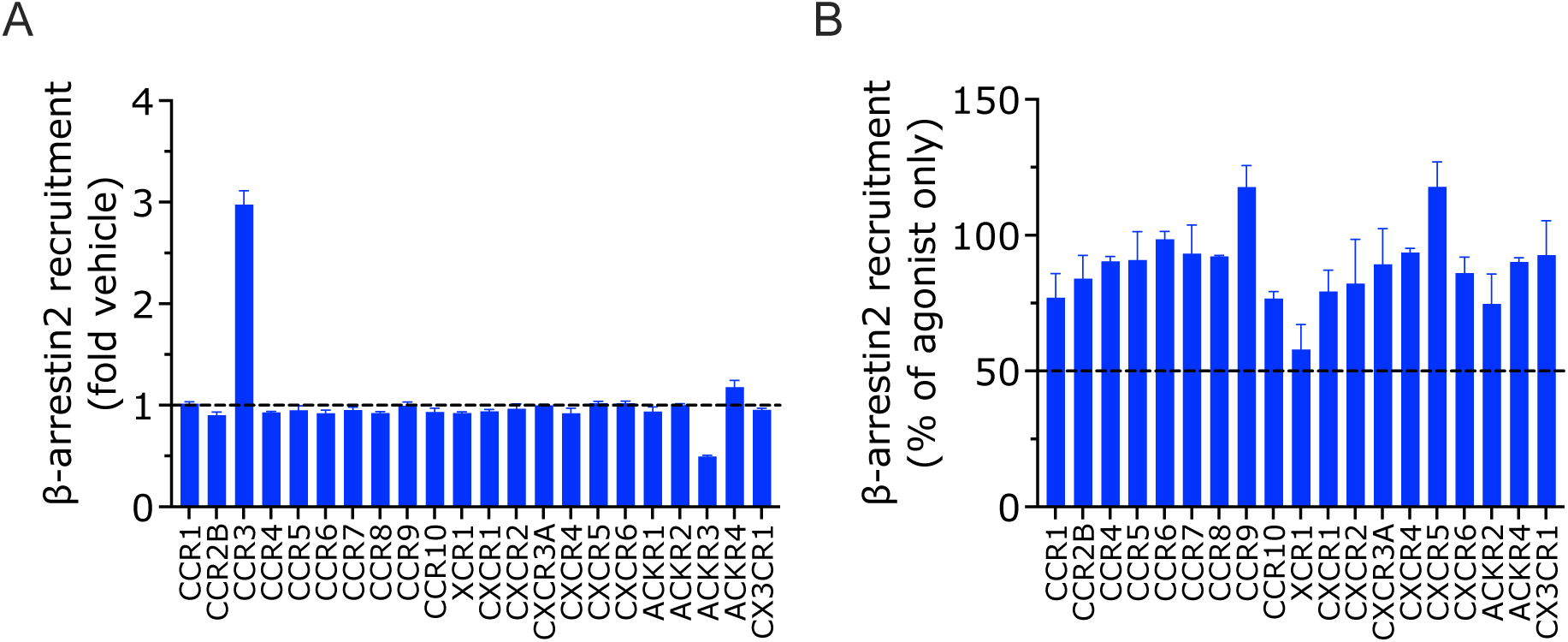
Modulation of a panel of chemokine receptors by VUF16840. Β-arrestin2 recruitment towards a full panel of 19 chemokine receptor was measured using a NLuc complementation assay in HEK293T cells. In panel **A,** VUF16840 (1 µM) mediated effect on β-arrestin2 recruitment is depicted, whereas in panel **B** VUF16840 (1 µM) was co-incubated with a receptor specific chemokine. All values depict the mean ± SD of ≥ 3 experiments.

In an antagonist-mode assay, VUF16840 was co-incubated with the respective endogenous agonist and in this set-up it did not show extensive inhibition (>50%) of any of the tested chemokine receptors (**Fig. 4B**). These data confirm that VUF16840 is at least 30-fold specific over CCR3 and shows an even larger selectivity over other tested chemokine receptors. It is anticipated that VUF16840 only modulates ACKR3 and CCR3 at pharmacologically relevant concentrations.

### VUF16840 inhibits constitutive ACKR3 signaling and trafficking events

Next, it was explored whether VUF16840 could also affect a number of additional ACKR3 signaling or trafficking events. Both β-arrestin1 and GRK2 are known to be recruited to hACKR3 upon activation^10,23,45,^ and phosphorylation of the C-terminus is found to be crucial for ACKR3 internalization and β-arrestin1/2 recruitment^10,47,48^, mediated at least in part by GRK2^10,47^. Additionally, clathrin A is a structural protein involved in the formation of clathrin-coated vesicles mediating receptor packaging and internalization, whereas Rab5a is a protein highly expressed in early endosomes ^32,49^. In order to explore how VUF16840 affects some of the events related to ACKR3 trafficking, BRET sensors in transiently transfected HEK293T cells were used to explore ACKR3 interaction with the following intracellular proteins: β-arrestin1, β-arrestin2, GRK2, Rab5a, and clathrin A (**Fig. 5**). The ACKR3 agonist VUF15485 induced recruitment of β-arrestin1 and GRK2 (**Fig 5A, B**) as well as translocation of ACKR3 towards early endosomes (**Fig 5C**), all with similar, high potencies (pEC_50_ values of 7.4 - 8.1; **Table S1**). Agonist-induced internalization of ACKR3 was further monitored by measuring the interaction between clathrin A and β-arrestin 2 using a NanoBiT-based protein-protein interaction assay (**Fig. 5D**). This interaction marks clathrin A-mediated endocytosis of GPCR bound β-arrestin2^32^ and was stimulated in a concentration-dependent manner by VUF15485 (pEC_50_ = 8.2 ± 0.4, mean ± S.D., n = 5; EC_50_ = 6 nM). In all these assay setups VUF16840 acted as an inverse agonist and inhibited the constitutive ACKR3-mediated responses with nanomolar potencies (pEC_50_ values of 8.0 - 8.8; **Table SI1**). In line with the effect of VUF16840 on β-arrestin2 recruitment to the ACKR3, VUF16840 could also fully inhibit (**Fig. 5E-H**) VUF15485-responses (pIC_50_ values 7.3 - 8.7; **Table S1**).

**Figure 5.**
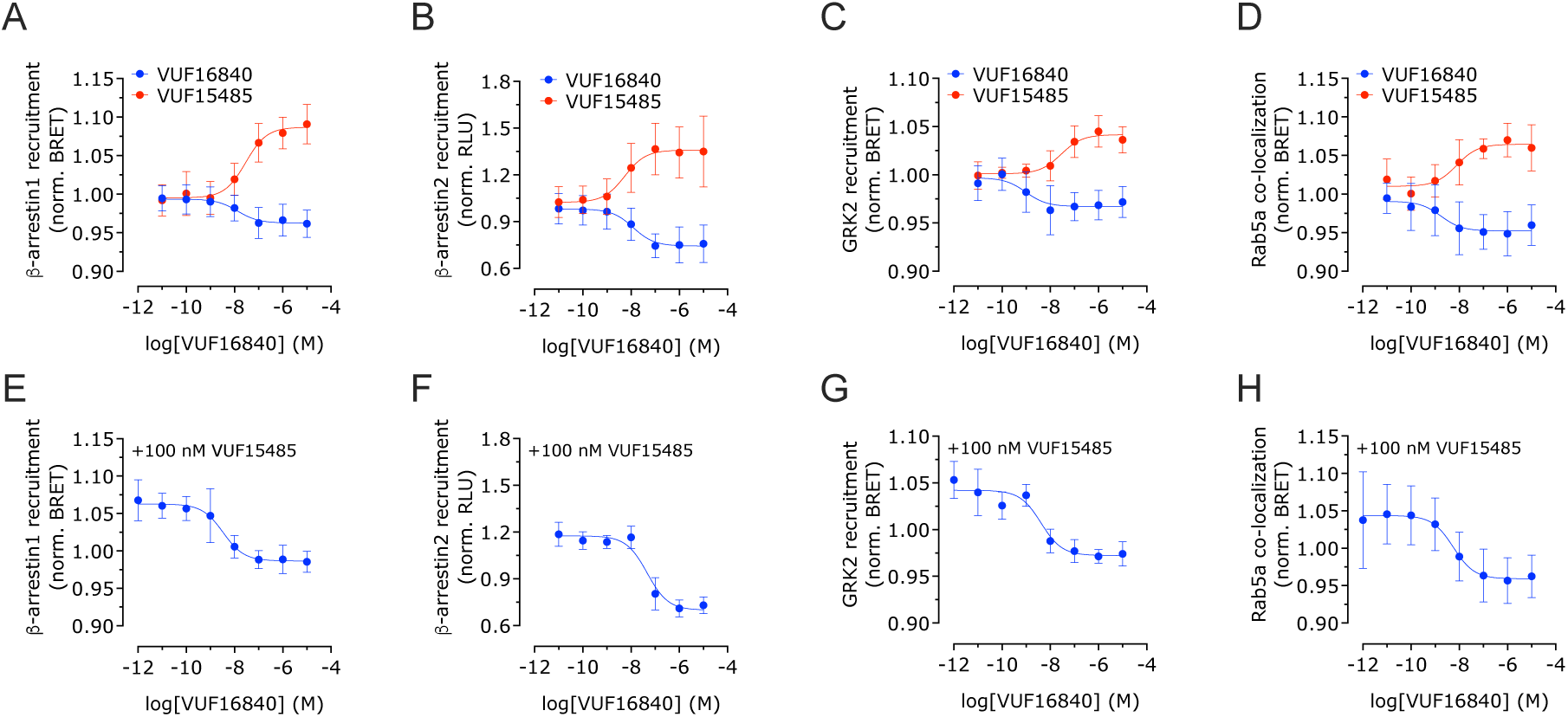
Inhibition of both constitutive as well as agonist-induced ACKR3 activity by VUF16840. HEK293T cells were stimulated with increasing concentrations of VUF15485 or VUF16840 only (**A-D**), or increasing concentrations of VUF16840 in the presence of 100 nM VUF15485 (**E-H**). The resulting proximity between ACKR3 and β-arrestin1 (**A, E**), GRK2 (**B, F**) or Rab5a (**C, G**) was determined by BRET. Furthermore, an interaction between clathrin A and β-arrestin2 was measured by NLuc complementation (**D, H**). Depicted data is normalized as fold-basal and represents the mean ± SD of ≥ 3 experiments with triplicate measurements per experiment.

ACKR3 cell surface expression was also studied by a bystander BRET method^37^ in which plasma membrane co-localization of hACK3-NLuc and membrane-targeted mVenus or mNeonGreen was measured (**Fig. 6**). As shown in **Fig. 6A**, CXCL12 exposure of hACKR3-NLuc expressing HEK293T cells resulted in a rapid reduction of bystander BRET signal from plasma membrane-targeted mNeonGreen, indicative hACKR3-NLuc internalization. Exposure of these cells to the inverse agonist VUF16840 increased the bystander BRET signals, indicative of receptor accumulation on the cell surface. In a similar setup with hCXCR4-NLuc, CXCL12 exposure also resulted in (sustained) CXCR4 internalization, but CXCR4 surface expression was not affected by VUF16840 (**Fig. 6B**). Moreover, bystander BRET between ACKR3-NLuc and plasma membrane localized mVenus-CAAX was concentration-dependently inhibited by CXCL12 (pIC_50_ = 9.1 ± 0.2, mean ± S.D., n = 3) and increased by VUF16840 (pIC_50_ = 9.1 ± 0.2, mean ± S.D., n = 3).

**Figure 6.**
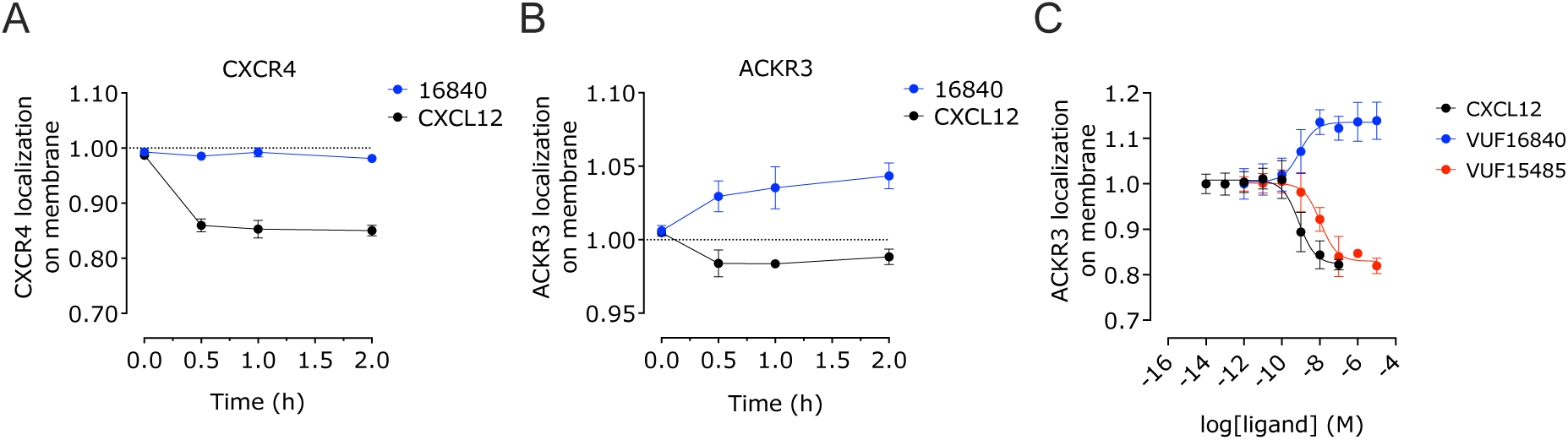
CXCL12 and VUF16840 differentially affect cell surface localization of hACKR3-NLuc. (**A**) Transiently transfected HEK293T cells, co-expressing hACKR3-NLuc and membrane-targeted mNeonGreen were incubated with 100 nM CXCL12 or 1 uM VUF16840. The resulting proximity between hACKR3-NLuc and was determined as bystander BRET. (**B**) The same assay was performed as in **A** for hCXCR4-NLuc. (**C**) Transiently transfected HEK293T cells, co-expressing hACKR3-NLuc and mVenus-CAAX were incubated for 15 min with increasing concentrations CXCL12 or VUF16840. Expression of hACKR3-NLuc at the cell surface was determined by the measurement of bystander BRET. Depicted data represents the mean ± SD of 3 experiments with triplicate measurements per experiment.

### The inverse agonist VUF16840 blocks CXCL12 uptake in ACKR3 expressing endothelial cells

Since VUF16840 inhibits CXCL12 binding to hACKR3 and affects also its constitutive internalization dynamics, we hypothesized that VUF16840 would also affect chemokine uptake via hACKR3. To evaluate the uptake of CXCL12 via ACKR3 internalization, we stably expressed hACKR3 in the endothelial cell line Human CMEC (D3) via retroviral transduction. As shown in **Fig. 7A**, the resulting D3-ACKR3+ cell line indeed internalized CXCL12-A647. After 60 min. incubation of D3-ACKR3+ cells with 3 nM CXCL12-AF647, significant intracellular accumulation of CXCL12-AF647 was observed. In line with the observed inhibition of ACKR3 internalization, co-incubation with VUF16840 resulted in dose-dependent inhibition of CXLC12-AF647 uptake (**Fig. 7B**). Following incubation of D3-ACKR3+ cells with 3 nM CXCL12-AF647 in the absence (DMSO) or presence of 100 nM VUF16840, cells were fixed and stained for actin and DNA with rhodamine-labelled phalloidin (red) and DAPI (blue), respectively (**Fig. 7C**). In the absence of VUF16840 CXCL12-A647 uptake could be observed (white dots), whereas VUF16840 treatment (100 nM) abolished the uptake of CXCL12-AF647 (**Fig. 7C**).

**Figure 7.**
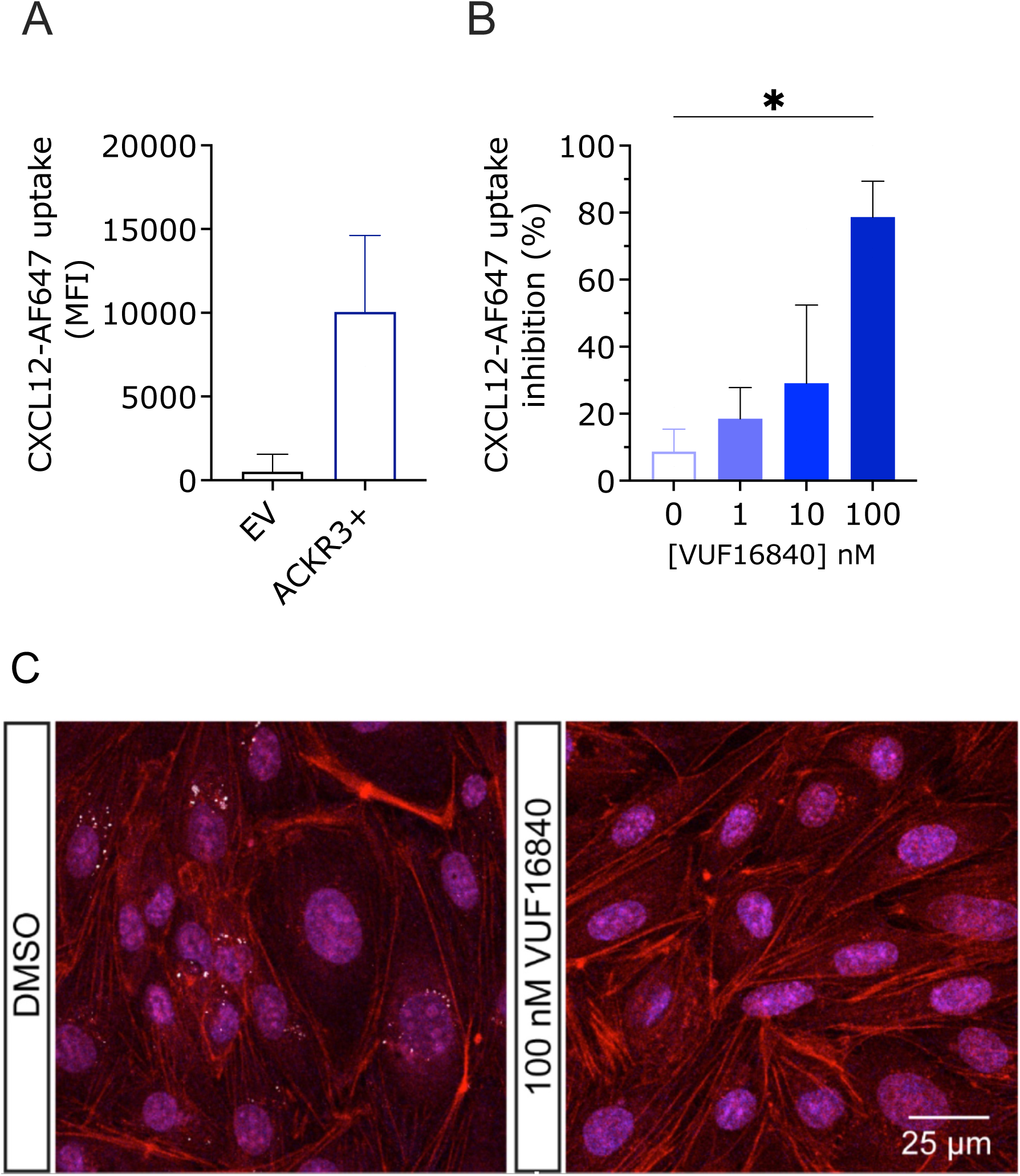
Fluorescent CXCL12-AF647 uptake by hACKR3-expressing D3 endothelial cells. (**A**) Stably transduced D3-ACKR3+ cells were incubated with 3 nM CXCL12-AF647 and fluorescent chemokine uptake in control (EV) and ACKR3+ cell was measured after 60 min. (**B**) Uptake of CXCL12-AF647 in D3-ACKR3+ cells in the absence or presence of increasing concentrations VUF16840. Depicted data represents the mean ± SD. (**C**) D3-ACKR3+ cells incubated with 3 nM CXCL12-AF647 in the absence (DMSO) or presence of 100 nM VUF16480, were fixed and stained with phalloidin (red, actin filaments) and DAPI (blue, nuclei) and imaged by confocal microscopy for CXCL12-AF647 uptake (white dots).

## Discussion

In this study we describe the pharmacological properties of a new high-affinity ACKR3 ligand, VUF16840. This compound combines nanomolar binding affinity for ACKR3 (**Fig 1B, Table S1**) with non-surmountable inhibition of chemokine-induced receptor activation, measured as ACKR3-β-arrestin2 interactions (**Fig 3**, **Table S1**). At high concentrations, VUF16840 completely prevents both CXCL11 and CXCL12 activation of ACKR3. A similar effect of VUF16840 is observed for ACKR3 inhibition following activation with the small molecule agonist VUF15485 (**Fig. S6**). Previously, most ACKR3 modulators with translational potential (e.g. CCX777 or CCX733) have been reported as agonists in ACKR3-dependent β-arrestin2 recruitment assays ^21,24^ and only recently ACKR3 antagonists were reported^50^. The chemical series of VUF16840 was originally identified within patent literature^25^ and during the course of this study compounds from the patent were published as non-surmountable ACKR3 antagonists^26,27^. Moreover, the selected clinical candidate from this series, ACT1004-1239, was reported to have therapeutic potential in animal models of MS and has entered clinical trials.^28–30^ Additionally, a different series of moderately active ACKR3 antagonists for the recruitment of β-arrestin2 was also reported by Pfizer recently.^50^ However, neither these ligands, nor ACT1004-1239 were shown to stabilize an inactive conformation of ACKR3, as is clearly the case for VUF16840 in our hands (**Fig 2A**, **2C**, **4A-D**). These data are corroborated by a preprint reporting on single molecule analysis of the conformational landscape of ACKR3 using VUF16840 as tool^51^. In the present study VUF16840 not only blocks chemokine-induced ACKR3 activation, but also inhibits constitutive recruitment of GRK2, β-arrestin1 and β-arrestin2 to ACKR3, next to constitutive interaction of ACKR3 with Rab5a and β-arrestin2 with clathrin A. The inverse agonism displayed by VUF16840 also affects ACKR3 expression at the plasma membrane (**Fig 6**). As reported previously^52^, ACKR3 traffics constitutively from the plasma membrane to Rab5a-expressing early endosomes (**Fig 4C**), most likely via clathrin A-coated pits (**Fig 4D**). Stimulation with agonists, such as CXCL12 can further increase the level of ACKR3 internalization, whereas exposure to the inverse agonist VUF16840 leads to its enhanced cell surface expression as measured by bystander BRET.

Inverse agonism by direct VUF16840 analogs, like ACT1004-1239, has so far not been reported^26^. This is not necessarily the consequence of structural and related pharmacological differences between those compounds and VUF16840, but potentially reflects differences in assay methodology, as the detection of constitutive GPCR signaling is known to rely on e.g. cellular background and GPCR expression levels^53,54^. Constitutive activity, the ability of GPCRs to signal in the absence of an external ligand, has emerged as a key feature influencing receptor physiology and pharmacology^53,54^. This phenomenon reflects the intrinsic capability of GPCRs to adopt active conformations spontaneously, enabling basal levels of signaling. Constitutive activity has been documented across various GPCR families, including (mutant) human chemokine receptors, such as CCR1^54^, CCR5^55^, CXCR3^56^ or CXCR4^57^. Consequently, several drug molecules targeting these receptors have been shown to act as inverse agonists^54–57^. A number of viral chemokine GPCRs (e.g. ORF74, US28) have previously also been shown to display a very high level of constitutive activity.^58^ For both CCR1 and the viral GPCR US28 the high level of constitutive signaling has been linked to increased levels of intracellular expression of the GPCR protein and internalization via β-arrestin-mediated mechanisms^54,58^, as in the case of ACKR3.

The constitutive recycling of ACKR3, similarly to the recycling of viral receptor US28^58,59^, has been linked to cellular uptake of chemokine ligands^49^. ACKR3 has therefore been hypothesized to act as a “chemokine sink” and e.g. indirectly affecting CXCR4 signaling^9,10,47,49,60^. In line with VUF16840 inhibiting constitutive recycling of ACKR3, the inverse agonist also completely blocks the *in vitro* uptake of fluorescent CXCL12 in endothelial cells stably overexpressing ACKR3. In addition, *in vivo* application of ACT1004-1239 has been shown to increase plasma and brain CXCL12 levels in mice^28,30^ and plasma levels in humans^61^, highlighting a potential relevant therapeutic effect of blocking the constitutive internalization of ACKR3.

In a selectivity screen with a wide panel of chemokine receptors, VUF16840 is shown to selectively interact with ACKR3 over most other chemokine receptors (**Fig 3**). The only observed off-target effect for VUF16840 was moderate agonistic activity on the CC chemokine receptor CCR3 (pEC_50_ 6.7). Despite a large difference in EC_50_ concentrations (> 30-fold selectivity), the cross-reactivity of VUF16840 at CCR3 warrants caution in complex *in vivo* models and e.g. the application of commercially available CCR3 antagonists (e.g. SB297006) to offset potential CCR3-activation by the VUF16840.

To understand the mechanism of inverse agonism and the non-competitive inhibition of agonist responses at the ACKR3 by VUF16840, it would be of great interest to understand the binding mechanism that is required to inactivate ACKR3. In a related study we recently combined hydrogen/deuterium exchange mass spectrometry, site-directed mutagenesis, and molecular dynamics simulations to show that VUF16840 and VUF15485 seem to largely occupy similar but distinct binding pockets of ACKR3^62^. Since several cryo-EM structures have in the meantime been resolved for peptide and small-molecule agonist-ACKR3 complexes^63^ or a β-arrestin2 x-ray structure bound to the C-tail of ACKR3^64^, structural biology approaches might in the future shed more light on the actual binding pocket of inverse agonists, like VUF16840.

In conclusion, our findings highlight VUF16840 as a potent and selective inverse agonist of ACKR3 with the ability to block both constitutive and ligand-induced receptor activation, internalization, and CXCL12 uptake. By stabilizing an inactive conformation, VUF16840 alters ACKR3’s trafficking dynamics, leading to enhanced plasma membrane localization and inhibition of its “chemokine sink” function. The compound’s selectivity for ACKR3, alongside its inverse agonistic properties, underscores its utility as a valuable tool for studying ACKR3 biology and pharmacology. Future research into the structural mechanisms underlying inverse agonism at ACKR3, including the precise binding interactions of VUF16840, will be instrumental in advancing therapeutic strategies targeting this atypical receptor in pathological conditions.

## Acknowledgements

This study was supported by the Health Holland TKI grant PROREMISE and the Luxembourg Institute of Health (LIH) through the NanoLux Platform, Luxembourg National Research Fund (grants INTER/FNRS 20/15084569 and CORE C23/BM/18068832) and the Fondation Cancer Luxembourg. Susane van der Pol and Bert (A.) J. van het Hof are acknowledged for technical assistance.

## Notes

### Competing Interest Statement

The authors have declared no competing interest.

