## Supplementary Information for "Inhibition of constitutive activity of the atypical chemokine receptor ACKR3 by the small-molecule inverse agonist VUF16840"

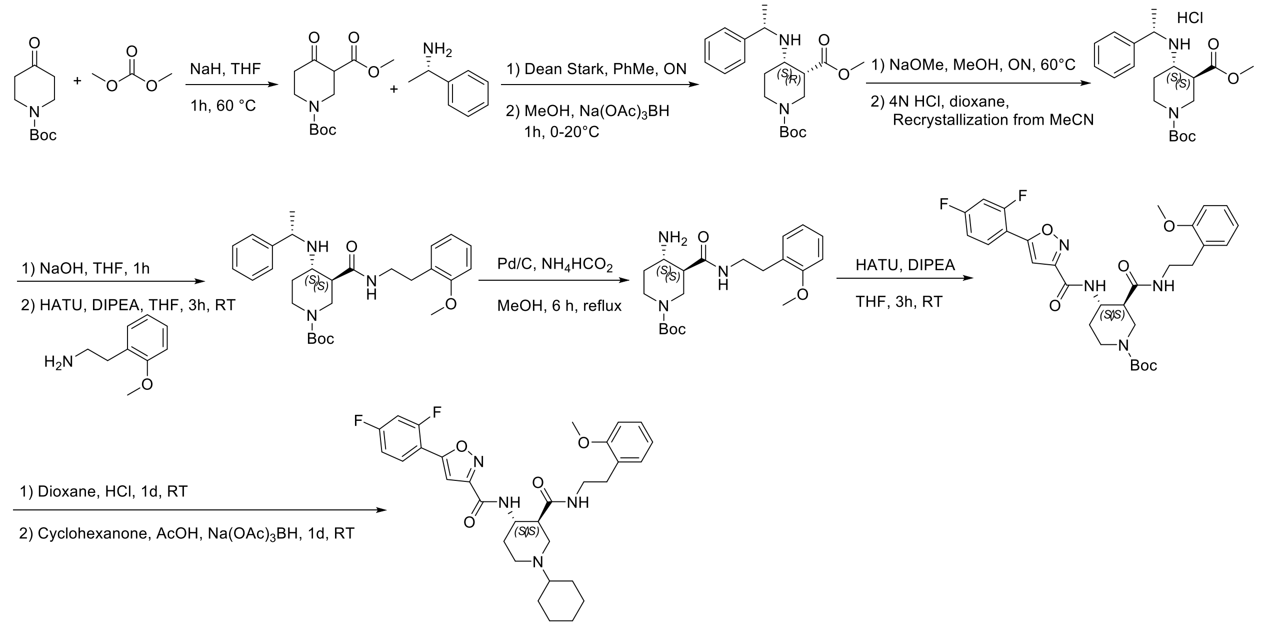


**Figure S1:** Synthetic scheme for VUF16840. This scheme was based with adaptations on a patent disclosure by Idorsia Pharmaceuticals. Details for the synthesis of a similar compound (clinical candidate ACT-1004-1239) were published during the course of our study[1,2]. ON=overnight. RT=room temperature.

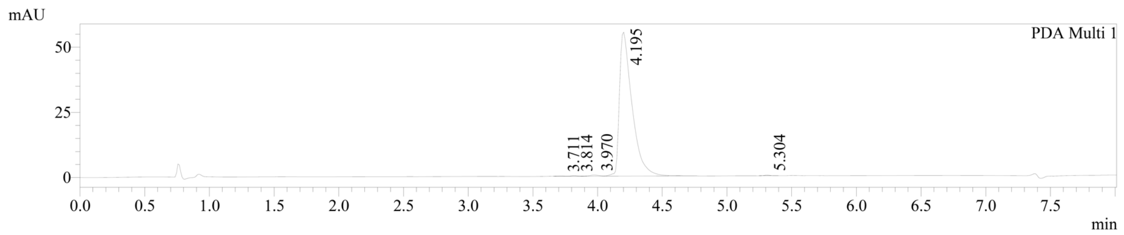


1 PDA Multi 1 / 254nm 4nm

| Peak# | Name | Ret. Time | Area | Area % |
|---|---|---|---|---|
| 1 | | 3.711 | 433 | 0.115 |
| 2 | | 3.814 | 854 | 0.226 |
| 3 | | 3.970 | 2179 | 0.577 |
| 4 | | 4.195 | 373461 | 98.901 |
| 5 | | 5.304 | 685 | 0.181 |

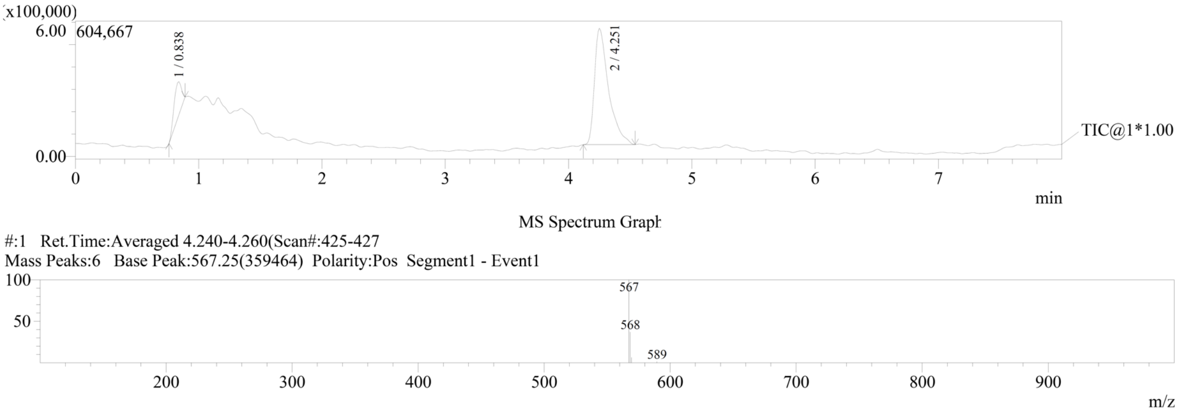


**Figure S2**: LC-MS analysis of VUF16840

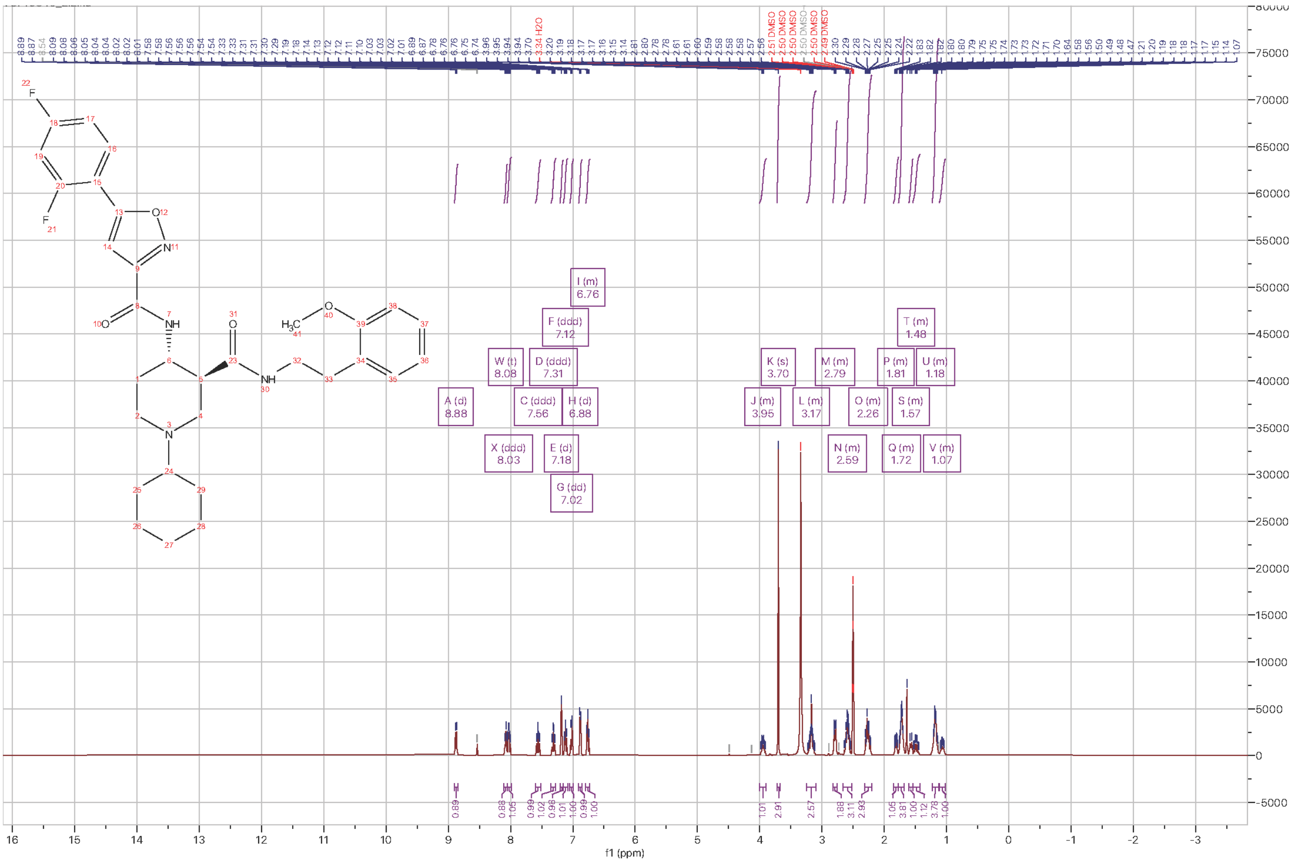


**Figure S3**: $^{1}$H NMR spectrum of VUF16840 (DMSO-d6)

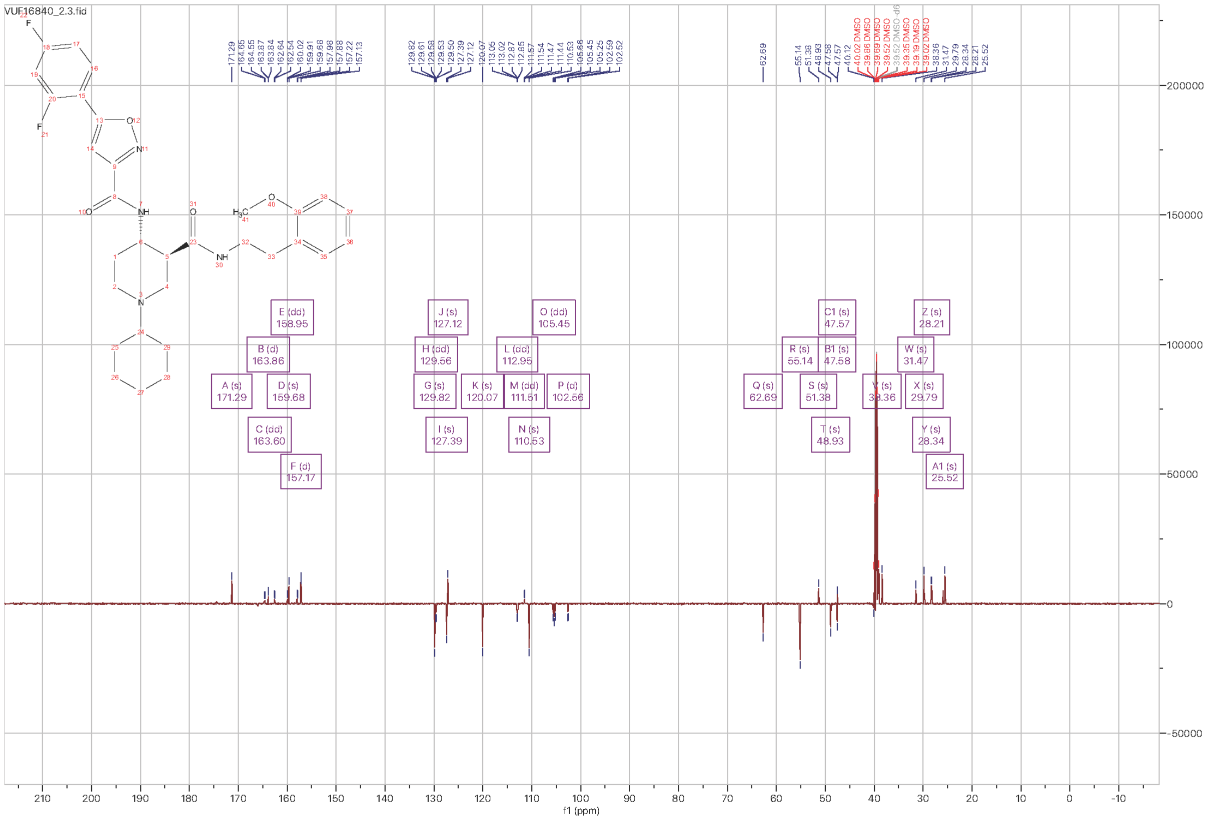


**Figure S4**: $^{13}$C NMR spectrum of VUF16840 (DMSO-d6)

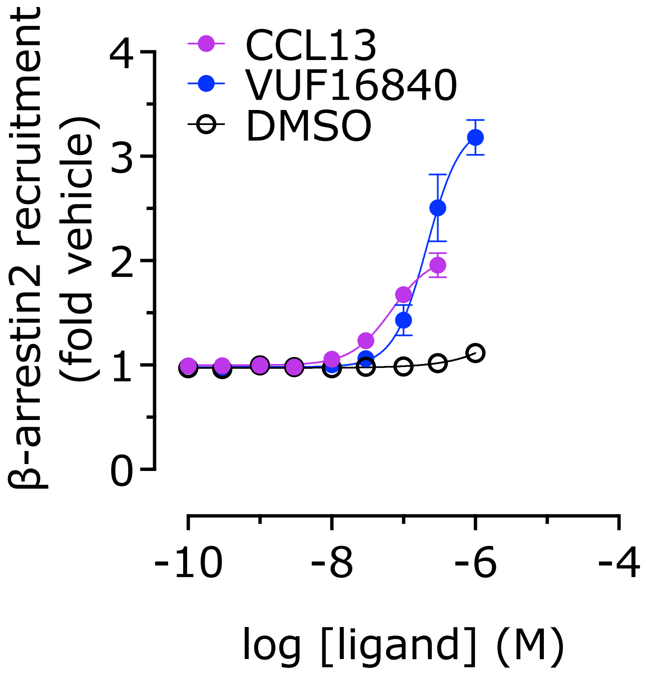


**Figure SI5 – Modulation of CCR3 receptor by VUF16840.** Recruitment of β-arrestin2 towards CCR3 chemokine receptor was measured using a NLuc complementation assay in HEK293T cells. The concentration-dependent activation of the CCR3 was determined for VUF16840 and the endogenous agonist CCL13. All values depict the mean ± SD of ≥ 3 experiments.

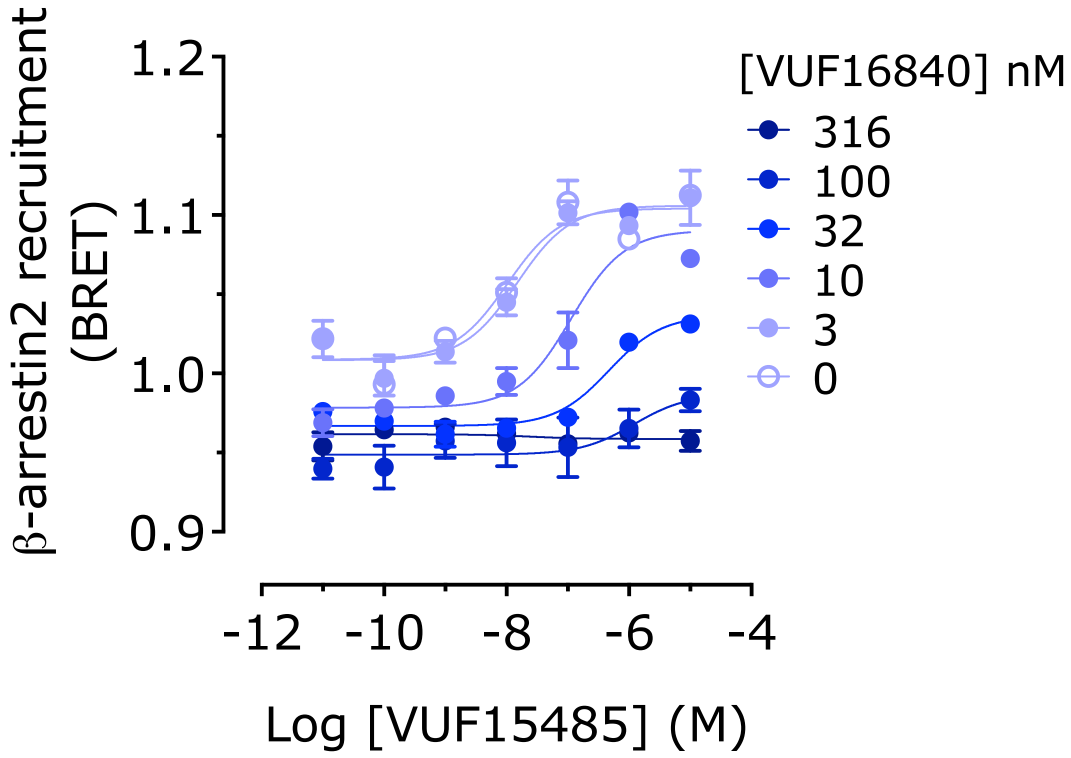


**Figure SI6 – Modulation of VUF15485-mediated ACKR3 activation by VUF16840.** Transfected HEK293T cells were stimulated with increasing concentrations VUF15485 in the absence or presence of increasing concentrations of VUF16840. All values depict the mean ± SD of ≥ 3 experiments.

**Table S1 –Ligand potencies to modulate the ACKR3.** The potencies of the inverse agonist VUF16840 and the agonists VUF15485, CXCL11 and CXCL12 to modulate ACKR3 are depicted as $pEC_{50}$ values. The inhibitory potency in which VUF16840 negates the effect of 100 nM of agonists VUF15485, CXCL11 and CXCL12 are depicted as $pIC_{50}$ values. All values depict the mean ± SD of (N) experiments.

| Mean ± SEM (N) | | | VUF16840 | VUF15485 | CXCL11 | CXCL12 |
|---|---|---|---|---|---|---|
| **β-arrestin2** | BRET | $pEC_{50}$ | 8.0 ± 0.1 (7)* | 7.9 ± 0.1 (7) | 8.2 ± 0.1 (3) | 8.5 ± 0.1 (6) |
| | | Inhibition by VUF16840 ($pIC_{50}$) | NA | 8.0 ± 0.3 (6) | 8.7 ± 0.2 (3) | 8.6 ± 0.2 (3) |
| | NanoBit | $pEC_{50}$ | 7.8 ± 0.1 (8)* | 7.8 ± 0.1 (7) | | |
| | | Inhibition by VUF16840 ($pIC_{50}$) | NA | 7.7 ± 0.0 (3) | | |
| **β-arrestin1** | BRET | $pEC_{50}$ | 8.3 ± 0.3 (6)* | 7.4 ± 0.2 (6) | 8.1 ± 0.1 (6) | 8.0 ± 0.1 (5) |
| | | Inhibition by VUF16840 ($pIC_{50}$) | NA | 8.7 ± 0.3 (4) | 8.6 ± 0.2 (4) | 8.3 ± 0.1 (4) |
| **Rab5a** | BRET | $pEC_{50}$ | 8.8 ± 0.3 (5)* | 8.1 ± 0.2 (4) | | |
| | | Inhibition by VUF16840 ($pIC_{50}$) | NA | 8.3 ± 0.2 (4) | | |
| **GRK2** | BRET | $pEC_{50}$ | 8.5 ± 0.3 (4)* | 7.6 ± 0.1 (3) | | |
| | | Inhibition by VUF16840 ($pIC_{50}$) | NA | 8.4 ± 0.1 (3) | | |
| **Clathrin A/ β-arrestin2 binding** | BRET | $pEC_{50}$ | 8.0 ± 0.3 (5)* | 8.2 ± 0.4 (5) | | |
| | | Inhibition by VUF16840 ($pIC_{50}$) | NA | 7.3 ± 0.2 (3) | | |

*$pEC_{50}$ value for VUF16840 as inverse agonist
